# The case for absolute ligand discrimination : modeling information processing and decision by immune T cells

**DOI:** 10.1101/019273

**Authors:** Paul François, Grégoire Altan-Bonnet

## 1. Introduction

Cells within the body are constantly bombarded with a large repertoire of molecules that must be dealt with as potential stimuli. Most of the time, these molecular inputs are measured by receptors at the surface of the cells. State of these receptors are thus informative on the outside world, and experimental and theoretical biophysicists [50] have extensively used information theory to estimate how much information (in the Shannon sense [45]) can be encoded at this stage. This information can be used and processed by the cells, and again transmission of information between Inputs and Outputs have been well studied (see *e.g.* in a developmental context work of Gregor and coworkers [22]).

Over longer time-scales, information processing eventually leads to a decision, that we define as a change in physiological macroscopic behavior or in the steady states of a gene regulatory network. For instance, in the well-studied example of bacterial chemotaxis, a cell might decide to switch behavior between tumbling or swimming [6]. Other examples include cellular commitment to a given fate in response to dynamical signaling pathways [9], or decision to take action in the case of immune responses, characterized by binary Erk phosphorylation, cytokine release, and cell proliferation [2].

There are obvious trade-offs between information content on the one hand, and cell decision (in particular its speed) on the other hand. For instance, it might not be desirable to optimize information collection if the environment changes too rapidly, as illustrated by the “info taxis” strategy [51]. In a context where decision needs to be taken rapidly, this can lead to discard or keep for later treatment some information content about the Input. For instance in the case of bacterial chemotaxis, cells take decisions by reading changes in the concentration of chemoattractants, and not absolute concentrations, and this can be well-explained by a Max/Min game theory model [8]. Additionally, information processing may be multi-tiered in order to retrieve different features of the input stimuli: rapid decision could discriminate between ligands of different quality, while slower decision could report the quantity of ligands.

In this review, we will focus on one specific cellular decision where similar considerations apply: the ability of T cells to discriminate very specifically between self and non-self ligands practically independently of their quantity. The functional significance of such ligand discrimination is quite obvious. “Recognize” a (potentially single) ligand as foreign, and a large set of responses is triggered to eradicate the pathogenic infection that generated this stimuli. Interact with (many present) self ligands as self and the T cell should remain quiescent to avoid auto-immune catastrophe.

This discrimination task is particularly daunting as T cells are constantly exposed to a large number of molecular stimuli at once. This issue of signaling pleiotropy is potentially a very generic problem in biology and we will coin the term “absolute discrimination” to describe it, Multitudes of receptors are indeed shared in examples as different as BMP signaling, olfaction, endocrine signaling, etc… or even in other immune contexts where fat-tailed distribution of antigen sequences is observed [41] and suggest strong overlapping signals.

The first part of this review will be devoted to a formal introduction to the problem of immune recognition by T cells, presenting current experimental understanding, past and present attempts to model this decision problem, and introducing the paradigm of adaptive sorting. In the second part we will introduce our current model for absolute immune discrimination, at the cellular scale. Finally, we will discuss how tools borrowed from statistical physics are needed to understand the higher level of processing in the immune system, at the cellular population scale.

## 2. Theoretical approaches for absolute ligand discrimination.

T cells probe their environment in search of potential foreign peptides. This is done via the interaction of their T cell receptors (TCR) with ligands (pMHC), presented by Antigen Presenting Cells. At a given time, these cells “present” a repertoire of oligopeptides (embedded within an MHC) that is representative of the current proteome (i.e. a mix of peptides from the self genome as well as a potential genome of the pathogen). The core function of T cells is to scan such repertoire and detect the presence of pathogen-derived ligands and respond, while not responding to self-derived ligands.

At the fundamental level, there does not exist any biochemical difference between self-derived pMHC and pathogen-derived pMHC, hence T cells must make a discrimination decision based on the biophysical differences between self and not-self.

Such decision must be, by essence, absolute in the sense that it must be determined by ligand quality, independently of ligand quantity. Absolute ligand discrimination is critical to enforce the functional goal of distinguishing self from not-self. In that context, decision has been shown to be logically all-or-none, via binary/bistable response in Erk phosphorylation [2, 36, 3] or in NFAT translocation to the nucleus. A very natural hypothesis would thus be that foreign ligands lead to specific allosteric modifications (conformational changes) at the level of T cells receptors, which would be an ideal way to confer extreme sensitivity and specificity to immune recognition. Molecular immunology has made a lot of progress in listing all components implicated in this early response, but could not find evidence for such a direct qualitative sensing in the general case. Indeed, differences between ligands rather appear to be of quantitative nature, explaining why mathematical and physical modeling must be called upon to address how continuous variation in ligand characteristics gets processed with absolute discrimination.

### 2.1. Insight from biophysics: the lifetime dogma of antigen discrimination

#### Antigen discrimination is set by the lifetime of the antigen/receptor complex

The exact molecular events associated with self/non-self ligand discrimination by T cells remain elusive. However, immunologists, structural biologists and biophysicists have made great progress to extract key parameters that physicists can build upon to tackle the issue of specific immune sensing (Figure 1.A).

**Figure 1:**
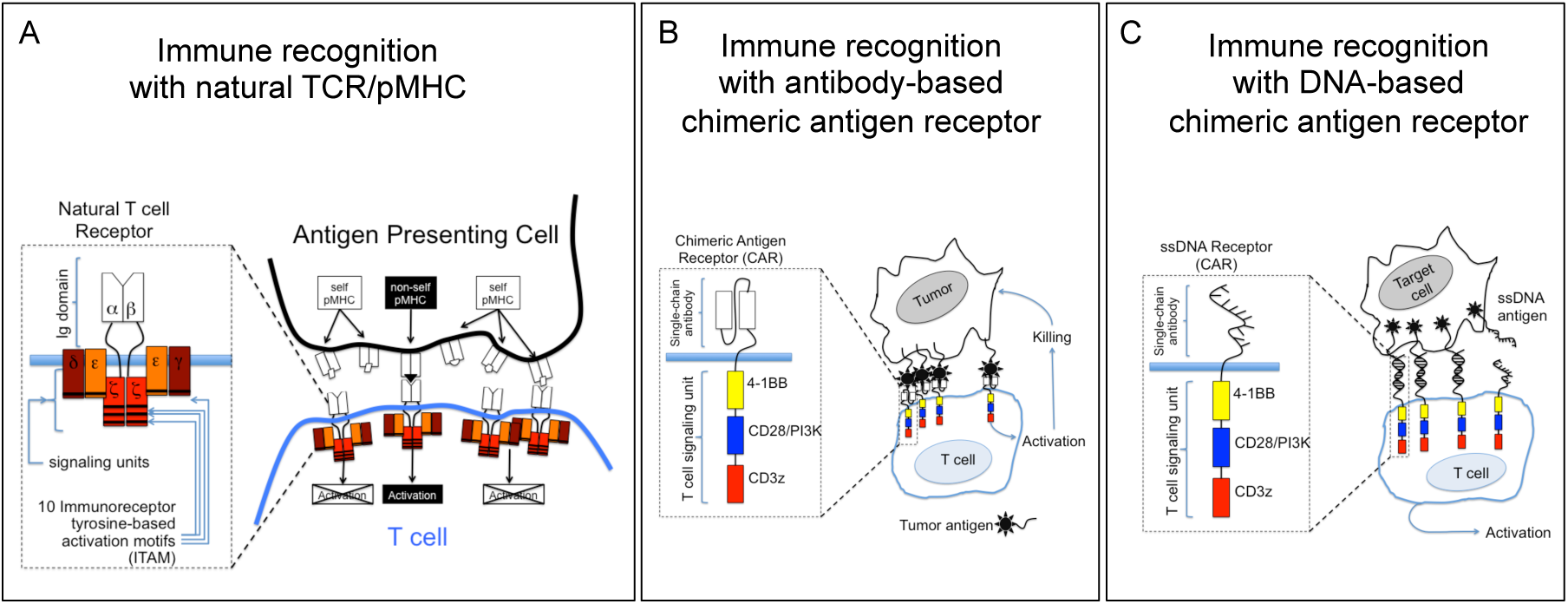
Three examples of immune recognition by T cells. **A.** The natural system of antigen discrimination by T lymphocytes relies on the T cell Receptor. It is composed of an extracellular a/b domains that interact with peptide-MHC complex on the surface of antigen-presenting cells, and 6 intracellular domains containing 10 Immunoreceptor Tyrosine-based Activation Motifs (ITAMs) that get phosphorylated upon engagement with non-self ligands, and trigger T cell activation. **B.** Recent developments in the field of immunotherapy introduced Chimeric Antigen Receptor (CAR): its extracellular domain is composed of a monomeric antibody that is specific for an antigen on the surface of the targeted tumor; its intracellular domain concatenates one ITAM-containing domain (z) and additional costimulatory domains (CD28 and 4-1BB) to induce robust T cell activation upon CAR engagement with its ligand. Such CAR combines the specificity of an antibody-based recognition, with robust signaling response. **C.** A DNA-based Chimeric Antigen Receptor was recently proposed by Ron Vale and coworkers [48]: its intracellular domain is based on the concatenation of signaling domains -similarly to the CAR described in B. Its recognition platform is composed of a single-stranded oligonucleotide, that recognizes another single-stranded oligonucleotide by sequence complementarity. Rather than relying on natural pMHC ligands whose biophysical characteristics are not tunable, Vale et al.’s design can be engineered to achieve variable lifetime for the ligand-receptor complex. Such ingenious experimental design will be critical to probe the sufficiency and limits of the lifetime dogma.

The first insight came in the 90s when researchers measured the biophysical characteristics of ligand-receptor interactions using purified proteins assayed *in vitro* for binding/debinding (*e.g.* detection by surface-plasmon resonance or by calorimetry). Qualitatively, experimental researchers documented that there exists a threshold of binding time (around 3-5s) so that for ligands with a lower binding time, T cells do not respond, while for ligands with higher binding time T cells do respond [21], thus realizing absolute discrimination based on binding. Such experimentally-derived rule (so-called “lifetime dogma” [15]) was well established by the turn of the millennium such that it became the springboard for many modeling efforts.

It should be immediately pointed out that like any dogma, this one is not absolute, and there are exceptions to the rule (see [15] for a discussion of some exceptions, or [13] for an experimental approach on cell populations). More recent measurements have been carried in the context of more complete biological systems (e.g. T cells reading their ligands on the surface of antigen presenting cells), with single ligand resolution: these brought about a correction on parameters for association and dissociation rates, but concurred qualitatively with the previously-acquired in vitro measurements. Recent work by Cheng Zhu and coworkers has challenged the lifetime dogma, using a force-measurement technique to resolve the dynamics of ligand–receptor engagement. Such exquisite technique leads to paradoxical results at first: strong ligands that trigger T cells were found to be weaker binder to the receptor, thus inverting the life-time dogma [28]. Subsequent studies tested how force loading on the ligand-receptor complex would alter its lifetime, and the more intuitive hierarchy of ligands was recovered, with better binders inducing better signaling responses [37]. Zhu and colleagues thus proposed a dynamics structural model, whereby agonist ligands induce a conformational change in the complex (so-called catch bonds) that triggers T cell activation. Alternatively, non-activating ligands (*e.g. self-derived peptide MHC*) would not induce such conformational change, would be released rapidly (so-called slip bond) and would fail to activate a significant signaling response. Hence, there would in the end be qualitative differences between activating and non-activating ligands. Still, such qualitative differences would mostly translate into quantitative differences for the ligand-receptor complex in its capacity to trigger signal transduction.

From the Physics point of view, one intriguing aspect of these measurements would be to add mechanical aspects to ligand discrimination. Understanding the forces associated with ligand-receptor interactions and the coupling with the mechanics of membrane deformation would be critical to account for the differential potency of ligands to activate T cells. Such quantitative models have been introduced [43], based on Ginzburg-Landau equations coupling the biochemistry of ligand-receptor interactions with the energetic cost of membrane deformation. Such models established that biochemical/mechanical coupling could be sufficient to physically sort membrane proteins on the T:APC cell interface, and generate a threshold of activation. Such physical models generated intriguing predictions that were subsequently validated experimentally: of note, it predicted that a family of ligands (with intermediate binding capacity) would abrogate the formation of so-called immunological synapse (a self-assembled bull-eye structure at the surface of T cells, where TCR aggregates at the center the synapse, and adhesion molecules occupy the periphery of synapse). In our context of immune recognition, one must point out that such synapse formation occurs downstream passed the initial signaling response associated with the ligand discrimination: it may constitute a reinforcing mechanism to anchor ligand discrimination over longer timescales, rather than the core cell-decision we are focusing on in this review. Recent models have explored how membrane stiffness influences effective binding times via suppradiffusive effects [1].

Two additional lines of work must be added to the biophysical conundrum of self/non-self ligand discrimination by T cells. First, in the field of immunotherapy, researchers have engineered T cells with synthetic chimeric-antigen receptors (CAR) whose extracellular domain is composed of an antibody recognizing a protein on the surface of tumors to be targeted (e.g. CD19 for B cell lymphoma), and whose intracellular domain is derived from signaling components of T cells (Figure 1.B): engagement of these receptors (with non-physiological ligands of surface antigens with very large lifetime) has been shown to be necessary and sufficient to activate T cells. In fact, examples of supra-physiological lifetimes for antigen/receptor complexes that lead to T cell activation were derived experimentally by in vitro evolution of the TCR/pMHC complex [25]. In the context of modeling early immune detection, this is relevant as the biophysics of ligand-receptor interaction are very different (with very large binding affinities), yet consistent with the lifetime dogma: antigen/receptor pairs with very strongly-held complexes, and very large lifetimes are indeed very stimulatory.

Another line of experimental evidence has recently been reinforcing the lifetime dogma. Markus Taylor & coworkers [48] engineered a new class of chimeric antigen receptors, whose extracellular recognition unit is composed of single-stranded DNA 1.C). Antigens for these T cells are composed of complementary single-strands of DNA (e.g. an oligomer of adenosines and cytokines, to avoid secondary structures). Hence immune detection in that context is highly tunable, quantifiable and easy to model: it is essentially the biophysics of DNA hybridization that drives the engagement of this artificial antigen receptor. In that context, Taylor et al. demonstrated that the association rates of these artificial receptor/ligand pair were essentially constant as it is limited by the nucleation of double stranding between two complementary DNA pairs. However, the dissociation rates are highly variable and essentially dominated by the free energy of double-strand formation. Hence, these DNA-based chimeric antigen receptor and ligands recapitulate the biophysical characteristics of ligand-receptor interaction in the natural immune detection context. Most strinkingly, Taylor et al. found that the lifetime dogma holds with a threshold of activation set around 3s for the lifetime of the antigen-receptor complex [48].

As of 2015, although the structural details of the early events in immune recognition by T cells remain elusive, the consensus around the lifetime dogma is thus holding and it is enabling physicists to build biochemically-explicit or phenotypic models of good biological significance [2, 20, 35]. It constitutes a rich paradigm for both theoretical and experimental biophysical considerations, and most of our discussion will be within this framework.

### 2.2. Setting the problem for physicists: what does absolute immune discrimination entails ?

In recent years, quantitative immunology has partially characterized the “phenotypic space” of T cells as a function of these parameters. A “golden triangle” characterizing immune response can be drawn [15], Figure 2 A. The first vertex of this triangle is *ligand specificity*, as encapsulated in the lifetime dogma described in the previous section: there exists an absolute discrimination threshold on ligand binding time, around 3-5 s.

**Figure 2:**
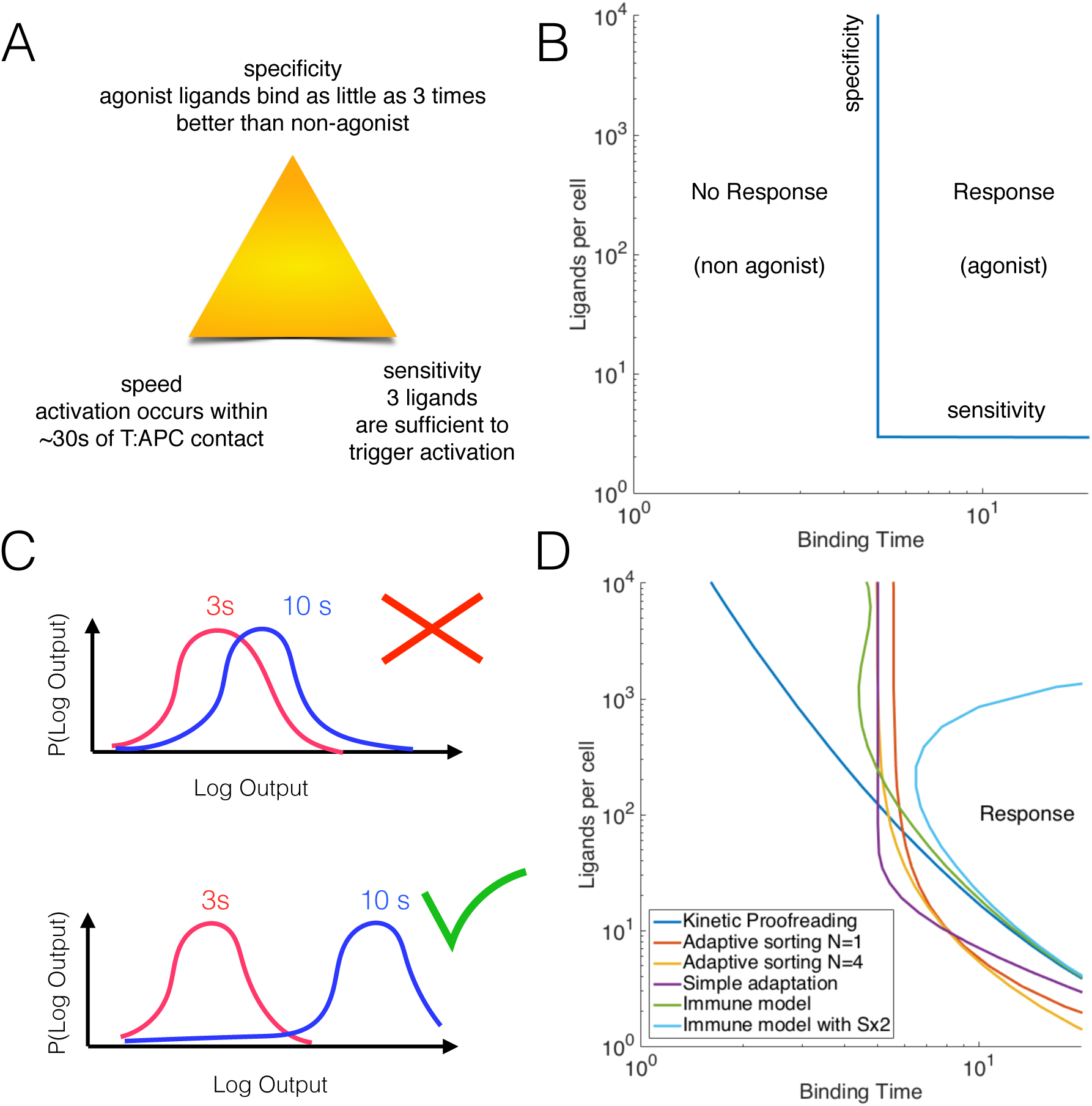
Absolute discrimination for physicists. A: Golden triangle for immune recognition. B: Idealized response line corresponding to the immune golden triangle. C: Two possible distributions for an Output variable directing immune decision, for networks exposed to random concentrations of ligands with identical binding times. In the top example, typical values for ligands with *τ* = 3 *s* and *τ* = 10 *s* overlap, so that it is not possible to discriminate between these ligands. In the bottom example, those distributions are well separated so that it is possible to choose a thresholding procedure on this Output to ensure absolute discrimination. D: Typical response lines for various models discussed in this review. Parameters were adjusted to have *τ*_*c*_ *∼* 5 *s*. Parameters for KPR and immune model from [20].

The second vertex of the triangle is *ligand sensitivity*. Minute amounts of ligand are able to trigger response. Actually, there are strong experimental evidence that one foreign ligand can trigger immune response [27], so that the physical limit of detection is reached biologically, a situation reminiscent of other famous examples such as photon sensing [5]. Such high sensitivity might be functionally critical as the immune system can “snip” a pathogenic infection before it has a chance to expand.

The last vertex of this triangle is *decision speed*. We know from experiments that immune decision at the single cell level is taken within a couple of minutes [55]. Note that this decision time depends quite strongly on ligand concentration [2] yet it is relatively fast to accommodate the limited time T cells spend scanning the surface of one antigen–presenting cells.

To reformulate this problem in a generic way, imagine a cell with a given set of identical receptors is suddenly exposed to *L* ligands, with identical binding time *τ*. We can plot in the (*L, τ*) plane a “response line”, characterizing the boundary between responding regions (”agonist” ligands) and non-responding ones (”non–agonist” ligands). The life-time dogma states that below some critical time *τ*_*c*_, there is no response, while above *τ*_*c*_ there is response. This defines a (vertical) “specificity” line in the (*L, τ*) plane. However, for high enough *τ*, it is clear that there should also be a minimum number of ligands triggering response: this defines a “sensitivity” line. It has been established experimentally that immune cells can trigger response when exposed to 1 to 3 agonist ligands. Thus the idealized lifetime dogma response line combines a vertical specificity line at *τ* = *τ*_*c*_, and a horizontal sensitivity line at *L ∼* 3 (Figure 2 B). A third dimension would be necessary to account for the third element of this triangle, speed, but is not drawn here.

### 2.3. Early attempts at modeling immune recognition: kinetic proofreading

Historically, McKeithan [40] was the first to propose a mechanistic model to underly the early events in immune recognition, accounting for such qualitative ligand recognition. The control of the quality of immune response by a single kinetic parameter *τ* is reminiscent of the famous Hopfield-Ninio kinetic-proofreading (KPR) paradigm, first proposed in the context of DNA replication and protein translation [26, 42]. In the immune context, McKeithan pointed out that subsequent to TCR-pMHC interactions, the receptor does internally go through several rounds of phosphorylation. He then assumed that, after unbinding of the pMHC ligand, receptor would be quickly dephosphorylated and the phosphorylation cascade would need to restart from “ground zero”. Calling *C*_*i*_ the ligand-receptor complex that has reached the *i*^*th*^ degree of phosphorylation in the cascade, *L* the ligand and *R* the receptor, simplified equations for a continuous model of this process are thus:

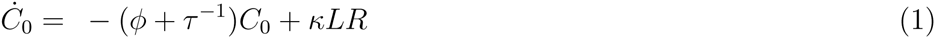

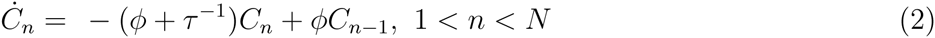

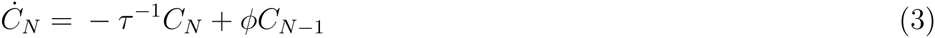

where *κ* is the association rate between ligand and receptor and *ϕ* the phosphorylation rate in the cascade. At steady state, assuming *R* is far from saturation and that *ϕ* << *τ*^−1^, one can easily derive that the last complex *C*_*N*_ thus has concentration scaling as *Lτ*^*N*+1^: this is the usual geometric amplification characteristic of kinetic proof–reading with *N* steps. It is thus clear that with only 2 steps, for the same initial ligand concentration, one can get up to 6 orders of magnitude in the difference of concentration of *C*_*N*_ for self ligands (*τ* = 0.1) vs agonist ligand (*τ* = 10). So if immune decision is taken via a downstream thresholding mechanism, physiological concentration of self ligands can not trigger immune responses even though one single agonist ligands theoretically can (see response line on Figure 2 D).

However, there are several quantitative shortcomings for a simple proofreading model. First, it is well known that to work efficiently, KPR needs to be very slow, which renders it incompatible with the observed fast response times of adaptive immune responses [2]. Second, the response line of KPR for any reasonable number of proofreading steps with realistic decision speed simply does not account for the observed specificity in terms of binding time *τ*, as illustrated on Figure 2 D. Finally, mixtures of ligand with different binding times will yield purely additive response to a kinetic proofreading mechanism, and thus would not account for more puzzling (but yet fundamental) aspects of immune response such as antagonism, where some not-reactive ligands can actually inhibit response of not self ligands [2].

Thus, one needs to augment the traditional KPR scheme with feedback regulation in order to be able to account for our “golden triangle” (specificity, sensitivity and speed) that characterizes the early events of T cell activation.

### 2.4. In silico evolution and adaptive sorting

The most counter-intuitive and puzzling property of immune response is its high specificity. The reason is that we would expect a priori that some shorter binding time could be “compensated” by higher ligand concentrations. More quantitatively, similar to the kinetic proofreading model, one would expect in general that any “output” *O* of a general signaling would behave as *O* = *f*(*L, τ*), where *f* is a monotonic function of both *L, τ*. But then how could we have a sharp process so that, on the response curve, a small decrease in *τ* (from agonist to self) leads to a change of *L* or several orders of magnitude ?

To answer this question, we turned to “in silico evolution” [33]. The general idea is to simulate a Darwinian process on the space of possible models and selecting for a desired phenotype (reviewed in [18]).

To define the phenotype to select, associated to absolute discrimination, we turn to mutual information, as a metric for the discrimination process. Assume a cell is exposed to one type of ligands with binding time *τ*, so that the probability *p*_*τ*_ (*L*) to observe a ligand concentration *L* is uniform on a log scale, within physiological concentration range. Consider an Output variable *O*. To each couple (*L, τ*) corresponds a distribution of output variable *O* characteristic of the signalling pathway, that we call *p*_*τ*_ (*O*|*L*). Since our problem for absolute ligand discrimination implies immune detection independently of ligand concentration, let us marginalize over all possible ligand concentrations and define a probability distribution for this Output associated to a binding time *τ* :

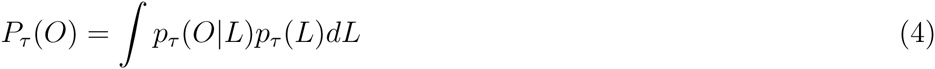

This probability distribution is then a pure function of the binding time of the ligand. Good ligand discrimination will be possible only if there is very little overlap between distributions corresponding to different *τs* (see Figure 2 C for an example of distributions with two different ligand types).

A practical way to use this for *in silico evolution* is to consider a situation where cells can be exposed to two different types of ligands (say *τ* = 3*s* and *τ* = 10*s* to fix ideas), with equal probability. Then, based on computed distributions such as the ones on Figure 2 C, we can easily define mutual information between the Output and *τ*, based on *P*_*τ*_ (*O*). A mutual information of 1 bit means that perfect discrimination is possible. Evolution *in silico* can then be used to optimize this mutual information. Using biochemical grammars inspired by early immune recognition, our procedure quickly converges to a very simple scheme described in [33], that we called “adaptive sorting” (Figure 3 A). Simplified equations for adaptive sorting are:

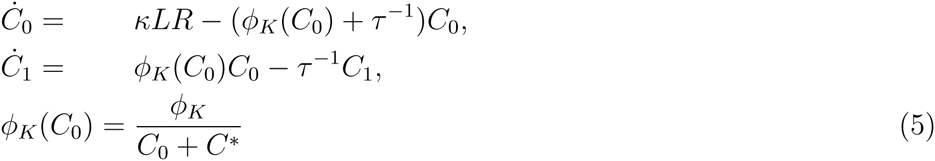

**Figure 3:**
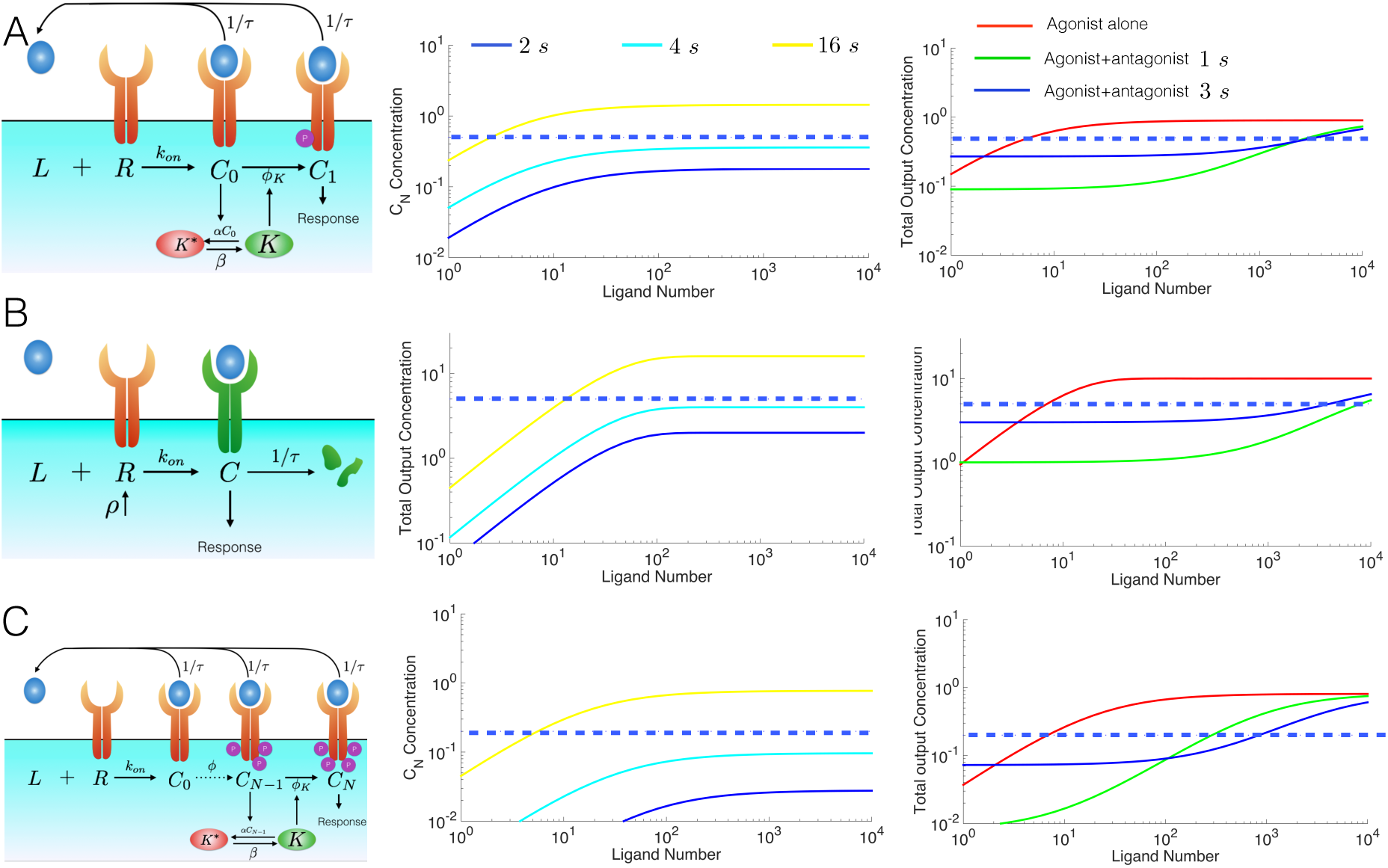
Examples of networks implementing the “adaptive sorting” paradigm. First column is network topology, second column is output concentration as a function of ligand for different values of *τ*, third column is output concentration as a function of agonist (*τ* = 10 *s*) ligand in presence of 10^4^ sub thresholds ligands, showing universality of antagonism. Dashed blue lines indicate thresholding for decision for deterministic systems used for simulations. A:Original Adaptive sorting evolved in [33]. B: Implementation of adaptive sorting based on the simplest two-variable model for biochemical adaptation evolved in [19] C: Elaborated adaptive sorting with more proofreading steps and reduced antagonism [33].

This network builds upon a minimum kinetic proofreading process by the addition of an incoherent feed-forward loop, where the first unphosphorylated complex in the cascade (*C*_0_) represses activity of the kinase *K* responsible for its own phosphorylation. Kinase *K* should diffuse rapidly inside the cell and be ’shared by all receptors, effectively coupling them, so that the total activity of the kinase *K* is a decreasing function *ϕ*_*K*_(*C*_0_) of *C*_0_. Steady state concentration of the phosphorylated complex down the cascade (*C*_1_) is then the product of *C*_0_ times the kinase activity *ϕ*_*K*_(*C*_0_), but those two terms are inversely correlated. As a consequence, *C*_0_ and thus *L* dependency in *C*_1_ can cancel out for some ranges of parameters, yielding an output

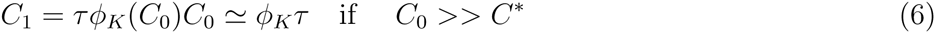

The later expression is independent from the amount of ligand presented, and then is a pure function of binding time thanks to the kinetic proofreading backbone as illustrated on Figure 3 A. Any thresholding process on *C*_1_ can thus efficiently discriminate between binding times. Response line of network is illustrated on Figure 2 D for a simple thresholding process on *C*_1_ (”Adaptive sorting N=1”), in close agreement with the “idealized” response from Figure 2 B.

### 2.5. Generalization of adaptive sorting to other pathways

The principle of negative feedforward interaction “compensating” to measure binding time of ligands can actually be generalized to any other biochemically adaptive network, to build generic networks able to perform absolute discrimination. For instance, let us consider the simplest possible adaptive network evolved *in silico* in [19] that is built from a simple ligand receptor interaction where receptor *R* is constantly produced/recycled and can only be titrated by ligand *L* :

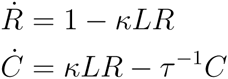

It is not unrealistic biologically to assume that parameter *τ* can also depend on the nature of Input ligand *L*, so that this network can use *C* steady state concentration *C*_*ss*_ = *τ* to measure ligand quality and thus eventually perform absolute discrimination (see Figure 3 B for an illustration and Figure 2 D for typical response line - “Simple Adaptation”.

More generally, any known adaptive network can be decomposed into one direct route sensing the input, and a “buffer” variable compensating the input variation pinning the output value back to a constant value (via a feedback or feedforward interaction) [19, 38]. If either the direct route or the “buffer” variable depends in some way on a biochemical parameter *k* characteristic of a ligand (on-rate, off-rate, etc…), then the steady state concentration of the output is independent of ligand concentration because of adaptation, but still sensitive to *k*. As such, “adaptive sorting” should be considered more as a general principle for absolute discrimination rather than a specific network topology.

### 2.6. Antagonism

Adaptive sorting is efficient to discriminate qualitatively between ligands with different binding times, and many different topologies perform similarly in terms of response line as illustrated on Figure 2 D. However, a more realistic immunological situation is that agonist ligands are presented simultaneously with many sub threshold (self) ligands. So cells not only need to discriminate agonists from self ligands, but should also detect agonists presented within many self ligands.

It turns out that the adaptive sorting schemes presented in Figure 3 A-B do not perform well: addition of many sub thresholds ligands considerably decrease output concentration for the same agonist concentration. In terms of response line of Figure 2 B, while specificity is conserved, addition of only a few self ligands actually shift drastically the sensitivity line upwards, meaning that the system loses sensitivity to minute concentrations of ligands, the second vertex of the golden triangle.

The fundamental reason for network in Figure 3 A is that interactions of self ligands with receptors eventually also leads to deactivation of kinase K, so that many more agonists ligands are required to trigger response. Such effect, called antagonism, actually appears to be a generic properties of systems performing absolute discrimination (P.F., in preparation). For instance, on the simple model of Figure 3 B, it is not difficult to see that steady state of total *C* activity for a mixture of two types of ligands *L*_1_, *L*_2_, with corresponding *τ*_2_ *< τ*_1_, is with transparent notations:

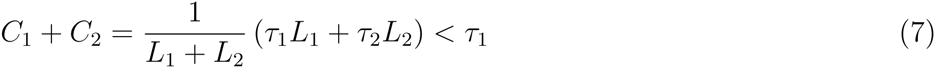

Since *τ*_1_ is the steady state of *C* when only ligands *L*_1_ are presented, it means than addition of ligands with lower *τ*_2_ effectively decreases total *C* and thus the effect of ligands *L*_1_. This can be especially important if some downstream decision is based on thresholding of total *C*: addition of subthreshold ligands could then push *C* below threshold, antagonizing decision. It is also clear that this effect is very strong: we basically need *L*_1_ *∼ L*_2_ for agonists to start overcoming the effect of strong antagonists.

As said above, it can be mathematically shown that antagonism is a necessary consequence of absolute discrimination (PF in preparation), and thus we can never completely get rid of it. However there is a simple way to minimize the range of binding times with strong antagonism in the model of Figure 3 A, by the addition of a short upstream proofreading cascade [33]. If repression of activity of the kinase *K* is made after a couple of proofreading steps, only ligands with already significantly high binding times will be able to deactivate the kinase. This will ensure that ligands with low binding times does not yield strong antagonistic effect while keeping intact the adaptive sorting property. This is illustrated on Figure 3 C: while response lines (Figure 2 D- “Adaptive sorting N=4”) and response to pure ligands is very similar to other models, the antagonistic effects are attenuated for sub thresholds ligands (third column of Figure 3).

While antagonism is reduced, there are however other trade-offs appearing in the system: for instance adding too many proofreading steps might reduce the final concentration of the output too much, which creates several downstream problems in terms of response times very similar to what happens in McKeithan’s KPR model (see Supplement of [34] for discussions of this effect).

### 2.7. Does adaptive sorting really satisfy the immune golden triangle ?

Adaptive sorting principle clearly satisfies by construction two constraints of the golden triangle: sensitivity and specificity. The third branch of the triangle, speed, is tightly related to another hidden problem of absolute detection: stochasticity of biochemical interactions. Considering the immune example, if a T cell is able to perform detection for ligand concentration as low as one ligand per cell, then one would expect a potentially deleterious sensitivity to intrinsic and extrinsic noise fluctuations.

A natural answer to this problem is to time-average response, but then it is not clear any more if a quick decision can be made. In the adaptive sorting mechanism, if we assume that *C*_1_ is activating a downstream “slow” output, it can be shown that indeed, intrinsic fluctuations can be averaged within a couple of minutes, which is then compatible with the experimentally observed immune decision time and the appropriate sensitivity and specificity of the TCR signalling pathway [33].

A related question is the minimum theoretical decision time for such process (”decision on the fly”). As said in introduction, a trade-off between accuracy and precision is expected (“exploit/explore”), and this clearly is an acute issue in an immune context. If decision takes too much time, a foreign ligand might be gone before proper response is activated. If accuracy is decreased, there is potential for auto-immune response. Exact results for this problem have been recently obtained by Siggia and Vergassola [46]. They study the problem of detection of change of composition of ligand mixtures. It turns out that Wald sequential probability test ratio on the log likelihood of a sequence of binding events can be used to take decision. Strikingly, simple phosphorylation networks reminiscent of network controlling early immune detection can naturally implement biological versions of this test [46]. An important aspect to optimize decision time is that decision here is made “dynamically”, while until now we have considered only networks performing decision at steady state.

## 3. Biologically realistic models for immune detection

As we have seen in previous section, generic solutions to the problem of absolute discrimination are now available within the simple adaptive sorting framework. But the natural question is to ask if biochemical reactions in actual T cells correspond to any of such Platonician view, and, if not, if they can be related to it in any ways.

Adaptive sorting elaborates on a small number of kinetic proofreading steps. It should thus be first pointed out that many (if not all) molecular components of the original McKeithan model [40] are indeed present in actual cells. Such “realistic” model explains its popularity and its use as a blueprint for many current models of immune decision [35]. The phosphorylation cascade would correspond to the known phosphorylation of “Internal Tyrosine Activation Motifs” (a.k.a. ITAM) containing chains in the TCR complex [29]. Rapid dephosphorylation upon unbinding would fit the “kinetic segregation” mechanism, specifying that generic phosphatases are segregated only upon ligand-receptor interaction [11]. Kinetic–proofreading–based models can account for other properties, such as potential dependency on the association rate of ligand with receptor, as insightfully discussed in [35].

### 3.1. Negative feedback and antagonism

Adaptive sorting based models are expected to include an extra negative element, *i)* buffering for ligand concentration to realize adaptation/absolute discrimination, and *ii)* associated with ligand antagonism. This has been indeed observed and characterized in seminal papers by Dittel *et al* [12], and Stefanova *et al* [47]. These papers establish the existence of a negative component in ligand detection via the Tyrosine-protein phosphatase non-receptor type 6 (PTPN6), also known as Src homology region 2 domain-containing phosphatase-1 (SHP-1), and its role in ligand antagonism. In details, Dittel *et al* quantified antagonism by measuring T cells responding to agonists in conjunction with increasing concentrations of antagonist ligands (which decreases the magnitude of immune response) in T cells endowed with two separate TCRs. Their measurement also suggested that antagonism is associated with SHP-1 association with TCR and the ensued dephosphorylation of ITAMs. Importantly, the antagonistic effect is “infectious” within the cells: recruitment by SHP-1 spreads from receptor bound to antagonistic ligands even to unbound receptors, suggesting a global coupling of receptors via SHP-1. Involvement of SHP-1 in antagonism is definitely established in [47]. They show in particular how moderate increase of SHP-1 activity gives several orders of magnitude increase of antagonism potency. Antagonistic ligands are also shown to recruit more rapidly SHP-1 compared to agonist ligands (which only recruit SHP-1 in a later time). They fianlly show that agonist ligand specifically trigger a positive feedback loop, mediated by ERK-1, and that ERK-1 specifically inhibit SHP-1 recruitment by the TCR, thus explaining the kinetic difference between ligands.

### 3.2. Modeling negative feedback, approximate adaptive sorting

The first model combining these different aspects was published by one of us (G.A–B) in collaboration with R.N Germain in 2005 [2]. This work, combined a kinetic proofreading backbone with a SHP-1 mediated feedback and an ERK-1 positive feedback. We included most known components of the system, including different co-receptors, kinases, and eventual phosphorylation cascade in a very complex mathematical framework including around 300 dynamical variables. This model succeeded in satisfying the previously described golden triangle as a modeling target. It also establishes a clear linear hierarchy for antagonism, where stronger antagonists are ligands with binding times just below critical threshold *τ*_*c*_. Most importantly, the full-blown model was tested with new experiments quantifying precisely antagonism strength and decision time of the network, and validating predictions from the model.

While the Altan-Bonnet Germain paper could explain many experimental features, it was not clear at that time if the full complexity of the model was required to understand the system. Can we this see the “biological wood emerge from the molecular tree” as nicely formulated by Gunawardena [23]? A first simplified model was proposed in 2008 by Lipniacki *et al* [36], but still contained around 40 variables, and was especially focusing on possible bistable properties of the system via ERK positive feedback loop. In 2013, we published in collaboration with G. Voisinne, E.D. Siggia & M. Vergassola a considerably simplified version of this model [20], focusing on the part of the decision network upstream of ERK. Schematic of the model is displayed in Figure 4 B. Like many other models, the backbone of this coarse-grained model relied on a kinetic proofreading backbone. Then, a simple negative feedback is included, in the form of the activation of a global phosphatase called *S* (corresponding biologically to SHP-1) by a single complex in the phosphorylation cascade.

**Figure 4:**
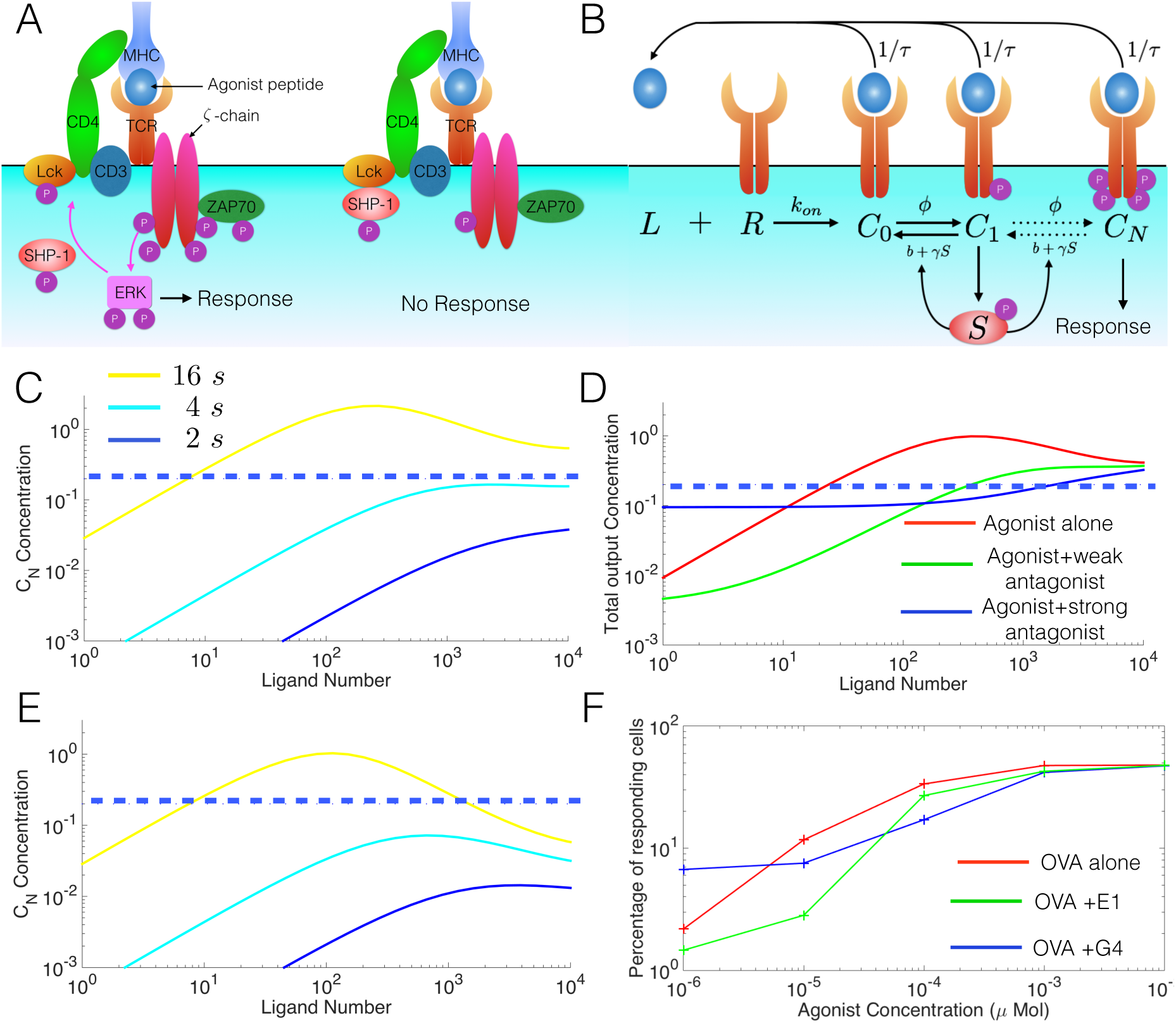
A simplified model for immune detection A: Sketch of some of the molecular players and their interactions. B: Simplified model based on kinetic proofreading combined with negative feedback C: *C*_*N*_ concentration as a function of ligand for different values of *τ* D: Total output concentration as a function of agonist (*τ* = 10 *s*) ligand in presence of 10^4^ sub thresholds ligands, showing antagonism. E: behaviour of the network when *S* total concentration is doubled, showing collapse at high ligand concentration. F: Experimental quantification of antagonism, reproduced from [20]. OVA is agonist, E1 is weak antagonist, G4 is strong antagonist.

**Figure 5:**
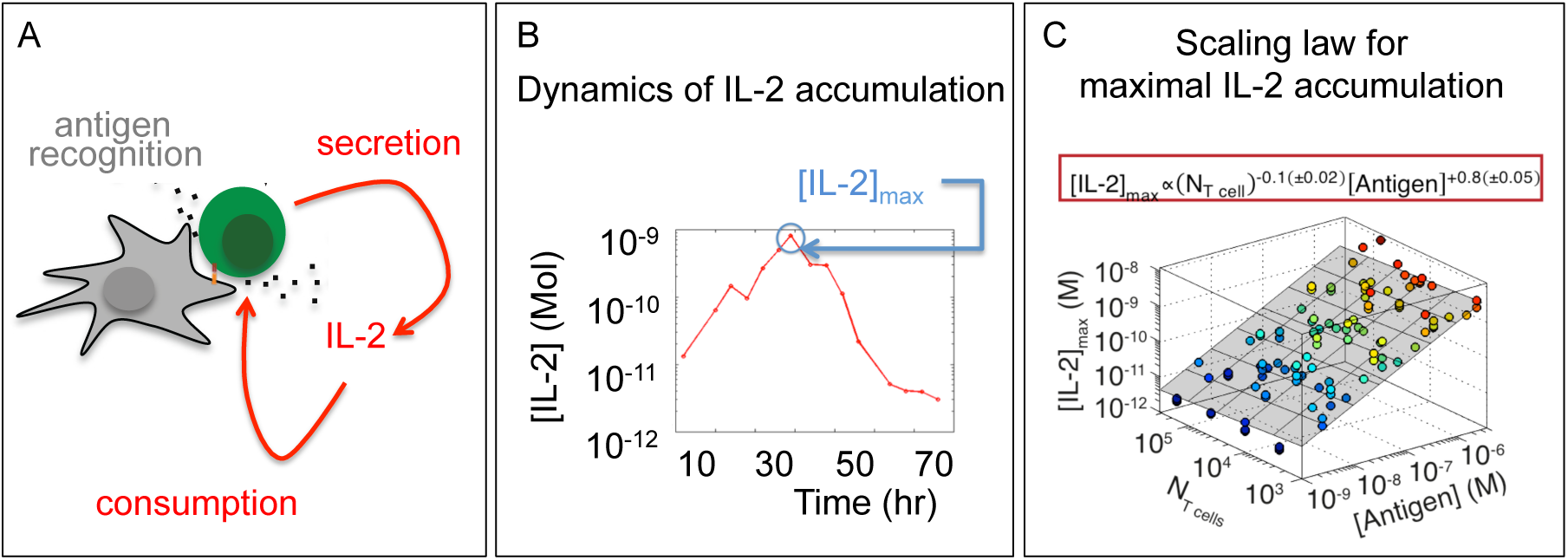
Immune recognition over long timescales may involve collective synchronization of T cell activation. While T cells perform an absolute (immune) discrimination at the individual cell level, their heterogenous expression of signaling components leads to unreliability in their threshold of activation. Moreover, experimental work on lymphocyte signaling demonstrated that individual T cells respond in an all-or-none (digital) such that their output contains limited information. Additional mechanisms must be at play to ??proofread?? individual immune recognition and/or generate plastic immune responses. Feedback regulations based on cytokine secretion and consumption constitute an efficient solution to the limitations of individual T cells [16, 24, 49, 54]. In particular, T cells can rely on such cell-to-cell communications to achieve higher levels of immune recognition. **B.** Tkach *et al.* found a surprising scaling law, whereby the maximum concentration ([*IL –* 2]_*max*_) of the IL-2 cytokine released by a population of T cells scales almost linearly with the amount of antigens that is present in the system, practically independently of the number of T cells present in the system. **C.** Such analog scaling at the population level was found to derive through a coherent feed-forward loop of Type 4, using the nomenclature introduced by U. Alon and coworkers [39].

This model essentially recapitulates all observations made in 2005 [2], satisfies the golden triangle, and is semi-analytic (and thus easier to study). Response line of the model is illustrated on Figure 2D (”Immune model”). Interestingly, this network actually performs an approximate adaptive sorting: it flattens out the concentration of the output *C*_*N*_ over several orders of magnitude of ligand concentration *L*, on different plateaus as a function of binding time *τ* as shown on Figure 4 C. Thus adaptive sorting appears to be the core principle of early immune detection, as could have been guessed in retrospect from first principles constraints evolved *in silico*. It is known that adaptation can be performed either via feedback or feedforward interactions, so that the network from Figure 4 B can be seen as a feedback version of the feedforward adaptive sorting network presented in Figure 3 C. Another difference with adaptive sorting as discussed before is that the same kinase and phosphatase is shared between all proofreading steps, while networks such as the ones displayed on Figure 3 A and C require one specific kinase for the step actually performing adaptation. So this model implements approximate adaptive sorting with a minimum set of unspecific kinases and phosphatases. Finally, only adaptive sorting models with several proofreading steps such as the one of Figure 3 C and Figure 4 give antagonistic properties similar to experimental data Figure 4 F [20]. Actually, it has been shown recently how such antagonism in adaptive sorting like models (including immune model from Figure 4) can help statistical decision in a fluctuating ligand environment [34], by buffering spurious ligand fluctuations.

We can also use this simplified model to simulate and even compute properties of mutants, where negative feedback does not saturate, which is relevant given the observed phenotypic variability (see next section). For instance, if we assume that phosphatase *S* (corresponding to SHP-1) is in excess, we have, for high concentration of agonist ligands, irrespective of binding times, a simple asymptotic relationship 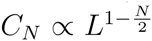 connecting ligand concentration *L* and *C*_*N*_, with *N* the number of proofreading step. Biologically, the highly non-linear feedback squashes response at high ligand concentration. There is a counter-intuitive prediction due to this behaviour: if feedback is strong enough, immune response at high agonist ligand concentration could thus disappear. Indeed, we tested and verified this prediction in cells with high level of SHP-1, see Figure 4 E (full stochastic treatment is presented in [20]). Furthermore, if SHP-1 level is increased above a couple a few-fold, negative feedback essentially dominates for all ligand concentration, and response is fully abolished, a “digital effect” first indeed experimentally observed in [17] and predicted with the Altan-Bonnet Germain model [2]

## 4. Ligand recognition by a population of lymphocytes: more is different?

### 4.1. Tackling cell-to-cell variability in immune responses: statistical physics to the rescue?

There are still many challenges to fully understand early immune response. In particular, once a cell has been activated, it appears that further processing occurs at the immune population level. Modeling the early events in immune detection potentially is very topical for statistical physics as it involves the accounting of cell-to-cell variabilities, modeling immune responses with distribution of cells, and the testing of their functional significance.

Starting with the seminal work of M. Elowitz *et al* [14], a subfield of biological physics has grown to address the emergence of cell-to-cell variability in biological systems. This corresponds to cases where an isogenic population of cells displays a distribution of phenotypes based on the heterogeneity of abundance of key regulatory proteins (receptors, kinases and phosphatases in the context of signal transduction network; transcription factors in the context of gene regulation). Such distributions are the physical consequences of stochasticity in the biochemical reactions and multiple sources of noise have been invoked (stochasticity of chemical reactions, epigenetic modulation of gene regulation etc.) *‡*.

First forays to address the functional significance of cell-to-cell variability in biological systems were focused on bacterial responses. From chemotaxis (the ability to orient motions in gradients of nutrients or chemokines) to competence the ability to acquire new genomic materials), researchers demonstrated that, indeed, varied levels of key proteins could map into varied phenotypes [30, 7]. Such observations were used at first as new quantitative constraints to validate biochemical models of biological regulation.

Concomitantly, these observations demonstrated that a “mean-field” measurement and model of a population of cells might have serious shortcomings when predicting global responses. One example where such cell-to-cell variability was found to be critical is in the study of bacterial antibiotic resistance. For example, Balaban and coworkers introduced a microfluidic device to track the proliferation and death of bacteria under antibiotic treatment [4, 44]. A subpopulaton of isogenic bacteria were found to resist to antibiotics, simply by being a different metabolic state compared to their sister cells at the time of exposure to antibiotics. Such process of distributed response based on distributed phenotype at stimulation time was described as an optimal strategy to tune responses in a fluctuating environment, by matching probability of phenotypic switching to probability of environmental changes [32]. Indeed, there is such a fundamental mismatch between the necessary response time (cells must exclude antibiotics on very short timescales) and the evolutionary constraints (it would take a large amount of time and a very low probability for cells to generate a solution to the problem of antibiotics resistance), that cells are better off diversifying their phenotype pre-emptively such that a solution is readily accessible when the antibiotic perturbation applies.

#### 4.1.1. Phenotypic variability of T cell ligand discrimination

Similar constraints are at play in the context of the immune response. There is a similar disconnect between the dynamics of biological problems at stake (eradicating a fast-replicating fast evolving pathogen vs. generating an adaptive immune response). In particular, recent measurements by the Jenkins & Davis lab have beautifully illustrated the number of constraints for a good immune response: the number of T cell clones that can recognize a specific antigen (e.g. flu peptide) is very small, with 10 to 100 clones per individual (mouse or human); on the other hand, lymphocyte can rapidly proliferate (by factors of 10^3^ to 10^6^) and relax back to low numbers for the memory pool. Such explosive proliferation is critical not only to match the challenge posed by fast-proliferating pathogens, but also to generate multiple cell fates to tackle pathogens invoking multiple escape mechanisms. Hence, rather than a deterministic adaptation of the immune response to the pathogenic challenge and optimization of detection at the cellular scale, one can conjecture that the immune system relies on some degree of statistical randomness in order to diversify responses within an isogenic population of lymphocytes.

The functional underpinnings of such cell-to-cell variability can be readily detected in the context of the early events of immune detection. Indeed, researchers have been using the vast panoply of antibodies to quantify protein and phospho-protein levels in cells, as well as the single-cell resolution of cytometry (using fluorescence-based or mass-spectrometry-based approaches), in order to quantify the cell-to-cell variability of responses to external stimuli. In a nutshell, if the abundance of protein X is limiting in the activation of Y into Y* (e.g. phosphorylation), then measuring and correlating X with Y* abundances at the single cell level will reveal heterogeneity of the response. Such cell-to-cell variability analysis has been carried out in the context of immune detection to demonstrate the sensitivity of ligand discrimination to varied levels of signaling components (e.g. CD8 and SHP-1) [17], as well as the sensitivity of T cells response to the cytokine IL-2 to varied levels of cytokine receptors (e.g. CD25, CD122, and CD132 a.k.a. IL-2R*α*, IL-2R*β* and *γ*_*C*_) [10]. Note that such parameter sensitivities were first predicted from the dynamical model of the signaling cascades at play, and CCVA validated these predictions quantitatively. Similarly, single-cell analysis has been carried out in the space of phosphor-proteins by the Pe’er & Nolan labs to quantify the strength of connections within the TCR signaling pathway [31]. There, analysis of single-cell measurements using overall, resolution of immune responses at the individual cell level has highlighted the large phenotypic variability in the signaling response of individual lymphocytes: such observations must then be interpreted functionally to map out how T cells diversify their response to optimize its detection capabilities.

#### 4.1.2. Deriving reliable immune responses from unreliable responses of individual cells

The cell-to-cell variability of lymphocyte response presents a challenge for our current cell-centric understanding of self/not–self discrimination in the immune system. Indeed, if each individual T cell makes a rapid, sharp yet utterly variable decision to respond to a ligand (see 4.1.1), one could anticipate many “mistakes” whereby T cells would respond to self tissues and trigger an auto–immune disorder.

We conjecture that it is the integration of the responses of individual T cells over longer timescales (*> hour*) that may correct for the “sloppiness” of individual T cell on short timescales (*≈* minutes). Such integration can be carried out through cell-cell communications *e.g.* through secretion and consumption of cytokines. In particular, K. Tkach *et al.* measured experimentally how much cytokine accumulates in the supernatant of T cells that were activated *in vitro* [49]. A surprising scaling law for this *IL –* 2_*max*_ capacity with the regards to the quantity of antigens #(*pMHC*) and the size of the T cell population is observed

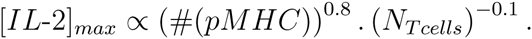

Hence, T cells are able to output cytokines in a near-linear scalable manner across four decades of #(*pMHC*): this is quite a striking observation when considering the limited dynamic range of individual T cell response and more generally of any biological system. Moreover, the independence of output with the number of T cells *N*_*T* *cells*_ in the system is remarkable considering that cytokine secretion is *a priori* an extensive variable: this finding implies that the overall capacity of the system (in terms of maximal cytokine concentration) is an intensive variable. Deriving an intensive output (*i.e.* an output that is independent from the size of the system) for a population of T cells has been observed in very related experiments [24]. Hart *et al.* measured the cell expansion of a population of CD4^+^ T cells after activation through their antigen receptor pathway. Hart *et al.*’s model focused on the cytokine IL-2 whose divergent function (cell proliferation and apoptosis) would explain such robust system-size-independent output.

Understanding mechanistically the emergence of such scaling laws implied revisiting our classical biochemical understanding of the IL-2 cytokine pathway. It was well established that T cells would essentially shut down their IL-2 production as soon as they secreted and built up a pool of shared cytokine. Biochemically, this implied that the IL-2 output of a population of T cells would have a ceiling of 10 *pM*, as the concentration of cytokine required to induce IL-2 signaling and subsequent shutdown to IL-2 secretion [53]. However, detailed quantitation of the biochemistry of the system unraveled a negative cross-talk from the antigen signaling response to the IL-2 response pathway [49]. When measuring the phosphorylation of the *ST AT* 5 transcription factor downstream of IL-2 sensing, Tkach *et al.* uncovered a surprising convolution between response to *pMHC* antigens and *IL*-2 cytokine:

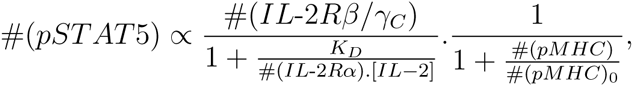

where #(*X*) represents the number of *X* within the cell. Such convolution between local and global responses (it i.e. *IL*-2 concentration and *pMHC* abundances respectively) was shown to be critical to regulate the *off* switch for *IL*-2 production. This is also important because *pST AT* 5 in turns regulate IL-2, which means that there is a feedback between the local stimulation (by *pMHC*) and the global readout (by *IL*-2) through the regulation of *ST AT* 5 phosphorylation. Overall, a combination of an incoherent feedforward loop was shown to be necessary and sufficient to explain the scaling law in the accumulation of *IL*-2 in the milieu.

Such initial attempt by Tkach *et al* [49] was based on detailed biochemical modeling with explicit biochemistry being implemented. Additional experiments revealed additional functional relevance for the secretion of IL-2, as a global regulator of self/non-self discrimination: Voisinne *et al.* probed the proliferation response of a polyclonal population of T cells, and demonstrated that weakly activated T cell clones could be co-opted by the activation of neighboring strongly activated T cell clones [52]. Similarly to previous studies [2, 16, 49], an experimentally-derived biochemically-explicit model of such multi scale integration was introduced to account for such lymphocyte co-optation. Overall, it is the addition of antigen signals (read locally through the TCR pathway) and cytokine signals (read globally through the IL-2 pathway) that decides T cell fate. Such observations expand over longer timescales (days) and larger timescales (lymph nodes) what individual T cell can contribute in terms of immune response. Future efforts will require phenomenological coarse-graining to allow better understanding of immune recognition at the level of the system.

Recent technical developments to monitor T cell responses at the individual cell level are enabling researchers to track the early events of immune activation, one cell at a time. Monitoring the differentiation of individual lymphocytes will certainly accelerate our study of the immune system, in particular when stochastic effects and phenotypic variability are necessary to explain the diversification in immune detection. Yet, in the context of the study of the immune system as a whole, it is the collective properties of the cells based on their cytokine communications and competition for antigen that shape the overall immune response. The future will certainly involve more statistical-physics based approaches that tackle the large number of lymphocytes and focus on emergent properties.

## 5. Wrapping up

To conclude, we presented an overview of some physics-inspired studies in immunology, focused on early recognition by T-Cells. Specifically, we discussed how the core problem of the adaptive immune response (the recognition of pathogen-derived antigens) must be addressed with quantitative models. Indeed, experimental results have now well established the biophysical underpinnings of self/not-self discrimination namely the fact that small increases in the lifetime of ligand-receptor complexes lead to large increases in ligand potency. Additionally, cells can respond very sensitively and with great speed. Quantitatively reconciling these three experimental aspects (so-called golden triangle) is a theoretical challenge with great conceptual and practical relevance (e.g. when fine-tuning T cell activation is required, as in cancer immunotherapies) We reviewed the current states of theoretical models, building upon the original kinetic proofreading scheme. *In silico* evolution of biochemical networks satisfying the golden triangle unraveled a minimal model that can reconcile all aspects of ligand discrimination. In particular, a proximal negative feedback (associated with the activation of a phosphatase) was found to be critically relevant to abrogate responses to self ligands (even in large quantities) while allowing responses to non-self ligands (even in small quantities). The functional pay-off of these models is to “predict” the existence of antagonism in immune recognition as well. Finally, we discussed how these models of ligand discrimination by T cells create new challenges in terms of understanding the phenotypic variability of isogenic populations of T cells, or in terms of accounting for the quantitative response to antigens when measuring T cell activation over long timescales. Similar collaborations between experimentalists and theoretical physicists will remain fruitful to expand our quantitative understanding of T cell activation to more complex issues in immunology (role of regulatory T cells, tuning of responsiveness according to inflammatory milieu etc.). More generally, we hope that these fundamental issues of immunology will spark the interest of statistical physicists, as the derivation and manipulation of large-scale immune response from the local activation of individual T cells remains poorly understood at the theoretical level.

## Footnotes

‡ note that, although physicists are fond to describe such variability as “noise”, based on their representation using a Langevin equation, this term remains confusing for most biologists because of its negative connotation: we elicit to use the more neutral term of cell-to-cell variability

## References

[1] Allard, J.F., Dushek, O., Coombs, D., van der Merwe, P.A.: Mechanical Modulation of Receptor-Ligand Interactions at Cell-Cell Interfaces. Biophysical Journal 102(6), 1265–1273 (2012)

[2] Altan-Bonnet, G., Germain, R.N.: Modeling T Cell Antigen Discrimination Based on Feedback Control of Digital ERK Responses. PLoS Biology 3(11), e356 (2005)

[3] Artyomov, M.N., Das, J., Kardar, M., Chakraborty, A.K.: Purely stochastic binary decisions in cell signaling models without underlying deterministic bistabilities. Proc Natl Acad Sci U S A 104(48), 18,958–18,963 (2007)

[4] Balaban, N.Q., Merrin, J., Chait, R., Kowalik, L., Leibler, S.: Bacterial persistence as a phenotypic switch. Science 305(5690), 1622–1625 (2004)

[5] Bialek, W.: Biophysics: searching for principles. Princeton University Press (2012)

[6] Block, S.M., Segall, J.E., Berg, H.C.: Adaptation kinetics in bacterial chemotaxis. Journal of bacteriology 154(1), 312–323 (1983)

[7] CaGatay, T., Turcotte, M., Elowitz, M.B., Garcia-Ojalvo, J., Suël, G.M.: Architecture-Dependent Noise Discriminates Functionally Analogous Differentiation Circuits. Cell 139(3), 512–522 (2009)

[8] Celani, A., Vergassola, M.: Bacterial strategies for chemotaxis response. Proceedings of the National Academy of Sciences of the United States of America 107(4), 1391–1396 (2010)

[9] Corson, F., Siggia, E.D.: Geometry, epistasis, and developmental patterning. Proc Natl Acad Sci U S A 109(15), 5568–5575 (2012)

[10] Cotari, J.W., Voisinne, G., Dar, O.E., Karabacak, V., Altan-Bonnet, G.: Cell-to-Cell Variability Analysis Dissects the Plasticity of Signaling of Common Chain Cytokines in T Cells. Science Signaling 6(266), ra17–ra17 (2013)

[11] Davis, S.J., van der Merwe, P.A.: The kinetic-segregation model: TCR triggering and beyond. Nature immunology 7(8), 803–809 (2006)

[12] Dittel, B.N., Stefanova, I., Germain, R.N., Janeway, C.A.: Cross-antagonism of a T cell clone expressing two distinct T cell receptors. Immunity 11(3), 289–298 (1999)

[13] Dushek, O., Aleksic, M., Wheeler, R.J., Zhang, H., Cordoba, S.P., Peng, Y.C., Chen, J.L., Cerundolo, V., Dong, T., Coombs, D., van der Merwe, P.A.: Antigen Potency and Maximal Efficacy Reveal a Mechanism of Efficient T Cell Activation. Science Signaling 4(176), ra39– ra39 (2011)

[14] Elowitz, M.B., Levine, A.J., Siggia, E.D., Swain, P.S.: Stochastic gene expression in a single cell. Science 297(5584), 1183–1186 (2002)

[15] Feinerman, O., Germain, R.N., Altan-Bonnet, G.: Quantitative challenges in understanding ligand discrimination by alphabeta T cells. Molecular immunology 45(3), 619–631 (2008)

[16] Feinerman, O., Jentsch, G., Tkach, K.E., Coward, J.W., Hathorn, M.M., Sneddon, M.W., Emonet, T., Smith, K.A., Altan-Bonnet, G.: Single-cell quantification of IL-2 response by effector and regulatory T cells reveals critical plasticity in immune response. Molecular Systems Biology 6 (2010)

[17] Feinerman, O., Veiga, J., Dorfman, J.R., Germain, R.N., Altan-Bonnet, G.: Variability and Robustness in T Cell Activation from Regulated Heterogeneity in Protein Levels. Science 321(5892), 1081–1084 (2008)

[18] François, P.: Evolving phenotypic networks in silico. Seminars in cell & developmental biology 35, 90–97 (2014)

[19] François, P., Siggia, E.D.: A case study of evolutionary computation of biochemical adaptation. Physical Biology 5(2), 26,009 (2008)

[20] François, P., Voisinne, G., Siggia, E.D., Altan-Bonnet, G., Vergassola, M.: Phenotypic model for early T-cell activation displaying sensitivity, specificity, and antagonism. Proc Natl Acad Sci U S A pp. 1–15 (2013)

[21] Gascoigne, N.R., Zal, T., Alam, S.M.: T-cell receptor binding kinetics in T-cell development and activation. Expert reviews in molecular medicine 2001, 1–17 (2001)

[22] Gregor, T., Tank, D.W., Wieschaus, E.F., Bialek, W.: Probing the limits to positional information. Cell 130(1), 153–164 (2007)

[23] Gunawardena, J.: Models in biology: ‘accurate descriptions of our pathetic thinking’. BMC biology 12(1), 29–29 (2013)

[24] Hart, Y., Reich-Zeliger, S., Antebi, Y.E., Zaretsky, I., Mayo, A.E., Alon, U., Friedman, N.: Paradoxical signaling by a secreted molecule leads to homeostasis of cell levels. Cell 158(5), 1022–1032 (2014)

[25] Holler, P.D., Holman, P.O., Shusta, E.V., O’Herrin, S., Wittrup, K.D., Kranz, D.M.: In vitro evolution of a T cell receptor with high affinity for peptide/MHC. Proc Natl Acad Sci U S A 97(10), 5387–5392 (2000)

[26] Hopfield, J.J.: Kinetic proofreading: a new mechanism for reducing errors in biosynthetic processes requiring high specificity. Proceedings of the National Academy of Sciences of the United States of America 71(10), 4135–4139 (1974)

[27] Huang, J., Brameshuber, M., Zeng, X., Xie, J., Li, Q.J., Chien, Y.h., Valitutti, S., Davis, M.M.: A single peptide-major histocompatibility complex ligand triggers digital cytokine secretion in CD4(+) T cells. Immunity 39(5), 846–857 (2013)

[28] Huang, J., Zarnitsyna, V.I., Liu, B., Edwards, L.J., Jiang, N., Evavold, B.D., Zhu, C.: The kinetics of two-dimensional TCR and pMHC interactions determine T-cell responsiveness. Nature 464(7290), 932–936 (2010)

[29] Kersh, E.N., Shaw, A.S., Allen, P.M.: Fidelity of T cell activation through multistep T cell receptor zeta phosphorylation. Science 281(5376), 572–575 (1998)

[30] Korobkova, E., Emonet, T., Vilar, J.M.G., Shimizu, T.S., Cluzel, P.: From molecular noise to behavioural variability in a single bacterium. Nat Cell Biol 428(6982), 574–578 (2004)

[31] Krishnaswamy, S., Spitzer, M.H., Mingueneau, M., Bendall, S.C., Litvin, O., Stone, E., Pe’er, D., Nolan, G.P.: Systems biology. Conditional density-based analysis of T cell signaling in single-cell data. Science 346(6213), 1250,689–1250,689 (2014)

[32] Kussell, E., Leibler, S.: Phenotypic diversity, population growth, and information in fluctuating environments. Science 309(5743), 2075–2078 (2005)

[33] Lalanne, J.B., François, P.: Principles of adaptive sorting revealed by in silico evolution. Physical Review Letters 110(21), 218,102 (2013)

[34] Lalanne, J.B., François, P.: Chemodetection in fluctuating environments: Receptor coupling, buffering, and antagonism. Proc Natl Acad Sci U S A (2015)

[35] Lever, M., Maini, P.K., van der Merwe, P.A., Dushek, O.: Phenotypic models of T cell activation. Nature Reviews Immunology 14(9), 619–629 (2014)

[36] Lipniacki, T., Hat, B., Faeder, J.R., Hlavacek, W.S.: Stochastic effects and bistability in T cell receptor signaling. Journal of Theoretical Biology 254(1), 110–122 (2008)

[37] Liu, B., Chen, W., Evavold, B.D., Zhu, C.: Accumulation of Dynamic Catch Bonds between TCR and Agonist Peptide-MHC Triggers T Cell Signaling. Cell 157(2), 357–368 (2014)

[38] Ma, W., Trusina, A., El-Samad, H., Lim, W.A., Tang, C.: Defining network topologies that can achieve biochemical adaptation. Cell 138(4), 760–773 (2009)

[39] Mangan, S., Alon, U.: Structure and function of the feed-forward loop network motif. Proceedings of the National Academy of Sciences of the United States of America 100(21), 11,980–11,985 (2003)

[40] McKeithan, T.W.: Kinetic Proofreading in T-Cell Receptor Signal-Transduction. Proceedings of the National Academy of Sciences of the United States of America 92(11), 5042–5046 (1995)

[41] Mora, T., Walczak, A.M., Bialek, W., Callan, C.G.: Maximum entropy models for antibody diversity. Proc Natl Acad Sci U S A 107(12), 5405–5410 (2010)

[42] Ninio, J.: Kinetic Amplification of Enzyme Discrimination. Biochimie 57(5), 587–595 (1975)

[43] Qi, S.Y., Groves, J.T., Chakraborty, A.K.: Synaptic pattern formation during immune recognition. Proceedings of the National Academy of Sciences 98(12), 6548–6553 (2001)

[44] Rotem, E., Loinger, A., Ronin, I., Levin-Reisman, I., Gabay, C., Shoresh, N., Biham, O., Balaban, N.Q.: Regulation of phenotypic variability by a threshold-based mechanism underlies bacterial persistence. Proceedings of the National Academy of Sciences of the United States of America 107(28), 12,541–12,546 (2010)

[45] Shannon, C.E.: A Mathematical Theory of Communication. Bell System Technical Journal 27(3), 379–423 (1948)

[46] Siggia, E.D., Vergassola, M.: Decisions on the fly in cellular sensory systems. Proc Natl Acad Sci U S A 110(39), E3704–12 (2013)

[47] Stefanová, I., Hemmer, B., Vergelli, M., Martin, R., Biddison, W.E., Germain, R.N.: TCR ligand discrimination is enforced by competing ERK positive and SHP-1 negative feedback pathways. Nature immunology 4(3), 248–254 (2003)

[48] Taylor, M., Jee, N., Gartner, Z., Mayor, S., Vale, R.D.: Stimulating T cel activation with DNA-based receptors and ligands. In: K. Symposia (ed.) T cells, regulation and effector function (2015)

[49] Tkach, K.E., Barik, D., Voisinne, G., Malandro, N., Hathorn, M.M., Cotari, J.W., Vogel, R., Merghoub, T., Wolchok, J., Krichevsky, O., Altan-Bonnet, G.: T cells translate individual, quantal activation into collective, analog cytokine responses via time-integrated feedbacks. elife 3, e01,944 (2014)

[50] Tkacik, G.G., Walczak, A.M.: Information transmission in genetic regulatory networks: a review. Journal of Physics, Condensed Matter 23(15), 153,102–153,102 (2011)

[51] Vergassola, M., Villermaux, E., Shraiman, B.I.: ‘Infotaxis’ as a strategy for searching without gradients. Nature 445(7126), 406–409 (2007)

[52] Voisinne, G., Nixon, G.B., Melbinger, A., Gasteiger, G., Vergassola, M., Altan-Bonnet, G.: T Cells Integrate Local and Global Cues to Discriminate between Structurally Similar Antigens. Cell reports 11(5), 1–12 (2015)

[53] Waysbort, N., Russ, D., Chain, B.M., Friedman, N.: Coupled IL-2-dependent extracellular feedbacks govern two distinct consecutive phases of CD4 T cell activation. J Immunol 191(12), 5822–5830 (2013)

[54] Youk, H., Lim, W.A.: Sending mixed messages for cell population control. Cell 158(5), 973–975 (2014)

[55] Zell, T., Khoruts, A., Ingulli, E., Bonnevier, J.L., Mueller, D.L., Jenkins, M.K.: Single-cell analysis of signal transduction in CD4 T cells stimulated by antigen in vivo. Proc Natl Acad Sci U S A 98(19), 10,805–10,810 (2001)

